# Acquired CRISPR spacers and rhamnose-glucose polysaccharide defects confer resistance to *Streptococcus mutans* phage ɸAPCM01

**DOI:** 10.1101/2025.05.05.652303

**Authors:** Lucas A. Wall, Daniel Wall

**Affiliations:** Department of Molecular Biology, University of Wyoming, Laramie, WY 82071, USA

**Keywords:** *Streptococcus mutans*, CRISR-Cas, bacteriophage, resistance, rhamnose-glucose polysaccharide, dental caries

## Abstract

*Streptococcus mutans* is a major cause of dental caries worldwide. Targeted therapeutic strategies to eradicate *S. mutans* include oral phage rinses. In this study, we investigated how phage resistance develops in *S. mutans*. As a model phage, we used ɸAPCM01, which is known to infect a serotype e strain. We isolated and sequenced the genomes of 15 spontaneous resistant mutants and found that 10 had acquired novel CRISPR spacers targeting the phage, with a total of 18 new spacers identified. Additionally, eight strains contained mutations in rhamnose-glucose polysaccharide (RGP) biosynthetic genes, three of which also acquired spacers. Only the *rgp* mutants exhibited defects in phage absorption, supporting the role of these cell surface glycans as the phage receptor. Mutations in *rgpF* and the newly identified gene *rgpX* led to severe cell division defects and impaired biofilm formation, the latter of which shared by the *rgpD* mutant. Thus, *rgp* mutations confer phage resistance but impose severe fitness costs, limiting pathogenic potential. Surprisingly, we found that ɸAPCM01 was capable of binding to and injecting its genome into UA159, a model serotype c strain. However, UA159 was resistant to infection due to an unknown post-entry defense mechanism. Consequently, ɸAPCM01 has the potential to infect both major serotypes associated with dental caries.

**Repositories:** The genome sequence of *Streptococcus mutans* DPC6143 was deposited at NCBI with the accession number NZ_CP172847.1.

## Introduction

Dental caries (tooth decay) is one of the most prevalent human diseases worldwide, resulting in billions of dollars in annual oral care (1). The Gram-positive pathogen *Streptococcus mutans* is most commonly associated with dental caries (2); therefore, targeted antimicrobial strategies against this bacterium represents a desirable treatment approach. Bacteriophage (phage) offer a promising strategy to specifically target and eradicate *S. mutans* while leaving oral commensal microbes unharmed (3). In light of this, seven *S. mutans*-specific phages have been isolated and initially characterized.

Of the seven *S. mutans* phages, six are lytic, and one is temperate. They all belong to the long-tailed Siphoviridae family. The phages M102, M102AD, ɸAPCM01 and SMHBZ8 are highly related, exhibiting 76−91% DNA sequence identity between them, with genome sizes ranging from 30−33 kb (4–7). The temperate phage ɸKSM96 genome is approximately 40 kb, and its host lysis and tail morphogenesis genes show up to 86% similarity to those of the previously mentioned phages, while other genes were more distant (8). Although phage e10 and f1 were not sequenced, DNA hybridization studies revealed they strongly hybridized to M102 (9). Consequently, all seven known phages share various degrees of relatedness suggesting common origins. Genomic and metagenomic analysis indicate the existence of additional *S. mutans* phage in nature (3, 10).

Along with the promise of developing new therapeutics to treat caries, phage serve as valuable tools to study the biology and pathology of *S. mutans*, including mechanisms of phage resistance for both applied and basic science purposes. For example, phage bind to cell surface receptors, and inactivating those receptors is a common mechanism of resistance (11). In turn, these mutations may result in changes in adhesion, biofilm, and/or fitness defects, among other phenotypes. In this context, *S. mutans*, like other *Streptococcus* species, possess active CRISPR-Cas systems, which provide adaptive immunity to phage infections by incorporating matching DNA spacers into their CRISPR array to target invading nucleic acids for degradation (12–14). These arrays also provide a chronological history of past phage encounters in ancestral lineages. *S. mutans* genomes typically contain one or two CRISPR-Cas systems, where many spacers match or nearly match known phages. Additionally, spontaneous resistant mutants to phage M102AD have been shown to acquire new spacers that matched the infecting phage (15). Interestingly, when these spacers or *cas* genes were deleted in all seven strains, they remained resistant to M102AD, indicating the presence of other unidentified mutations. Moreover, among dozens of resistant mutants that acquired new spacers, the majority were defective in phage adsorption, again suggesting additional mutations play a role in phage resistance.

To date, no study has characterized spontaneous phage-resistant mutants that map outside of CRISPR loci. However, targeted gene knockouts and allelic replacements of the rhamnose-glucose polysaccharide (RGP) biosynthetic pathway in a serotype c strain blocked infection and adsorption of phage M102, suggesting that cell surface RGPs serve as a receptor (16), which aligns with general observation that cell surface glycans frequently act as phage receptors in Gram-positive and -negative bacteria (11, 17, 18). In another study, an engineered missense mutation in the *S. mutans* methionine aminopeptidase resulted in a complete block in M102AD growth after adsorption, replication and protein expression had occurred (19). Together, these findings indicate that identifying spontaneous *S. mutans* mutations will provide insights about phage interactions.

In this study, we isolated, sequenced, and characterized 15 resistant mutants to ɸAPCM01, which infects a sereotype e *S. mutans* strain, DPC6143 (4). Genome sequencing identified newly acquired CRISPR spacers and gene mutations involved in resistance. Some of these mutations were located in RGP biosynthesis genes that reduced phage absorption. Additionally, *rgp* mutants exhibited defects in biofilm formation and cell division, and they readily clumped in solution. These findings and their significance are discussed.

## Methods

### Strains and culture conditions

Bacteriophage ɸAPCM01 was isolated from a subject’s saliva collected at the University College Cork Dental Hospital (4). The host strain DPC6143 (WT), a serotype e strain, originated from the Moorepark Food Research Centre collection in Ireland. The phage and host strain were kindly provided by Dr. Colin Hill. *S. mutans* DPC6143 was grown in Brain Heart Infusion (BHI) medium (Difco™ BHI Broth) at 37°C under aerobic conditions without shaking. *S. mutans* UA159 was kindly provided by Dr. Jacqueline Abranches and was similarly grown, except under anaerobic conditions using a BD GasPak™ EZ system. Phage ɸAPCM01 was propagated on DPC6143 in BHI medium supplemented with 10 mM CaCl₂.

### Mutant isolation

Two approaches were used to isolate phage-resistant mutants. In each case, the selected mutants were derived from independent overnight cultures of DPC6143 started from frozen stocks. In total, 25 cultures were screened.

Liquid lysate approach: Overnight cultures were grown, and 1 mL of culture was added to 5 mL of fresh BHI broth containing 10 mM CaCl₂ and approximately 10³ pfu of ɸAPCM01. The infected cultures were incubated at 37°C for one to four days. Afterward, 0.2 mL of the culture lysate was plated onto BHI plates and incubated for three days at 37°C to allow candidate phage-resistant colonies to form. Emerging colonies were picked and cross-streaked against a stock of ɸAPCM01. Resistant isolates were identified and further tested by serial dilutions (described below). Phage sensitivity was quantified, and resistant isolates were stored at −80°C for future use.

Plate lysate approach: Overnight cultures of DPC6143 were started from frozen stocks. A 1 mL aliquot from each overnight culture was added to 4 mL of 0.5% molten agar, 10 mM CaCl₂, and 100 µL of phage (10⁹ pfu/mL). This mixture was poured onto BHI (1% agar) plates to form a soft agar overlay. Plates were incubated at 37°C for five days to allow colonies to emerge. Colonies were selected, screened, and stored as described above. In all cases only one isolate from each batch was retained for genomic sequencing and further analysis.

### Phage sensitivity

Overnight cultures of DPC6143 and phage-resistant mutants were adjusted to an OD_600_ of 0.5 and 1 mL aliquot of culture was mixed with 4 mL of 0.5% molten BHI agar containing 10 mM CaCl₂ and overlaid on BHI plates. Tenfold serial dilutions of a phage stock (2×10⁸ pfu/mL) were spotted onto the *S. mutans* lawns and incubated for one day. Plaques were enumerated and compared between strains to calculate the efficiency of plating (EOP).

### SYBR Gold phage labeling fluorescent microscopy

SYBR Gold (Invitrogen) was diluted in TE buffer (pH 8.0) to prepare a 5,000× stock solution. It was then added to a phage stock (1×10⁹ pfu/ml) to a final concentration of 100× and incubated overnight at 4°C. After incubation, the labeled phage was purified by centrifugation using Amicon® Ultra 0.5 mL filter tubes for six min at 18,000 rpm. The phage were washed three times with buffer (20 mM Tris-HCl [pH 7.2], 10 mM NaCl, 20 mM MgSO₄) and centrifuged again for six min at 18,000 rpm. After discarding the wash, the phage were eluted with 100 µL buffer into a new collection tube, followed by filter tube inversion and centrifugation for three min at 15,000 rpm. Labeled phage were stored in the dark at 4°C until use.

To visualize phage, they were added to overnight *S. mutans* cultures in the presence of 2 mM CaCl₂ at approximately a 25:1 phage to cell ratio. For phage binding, samples were incubated for 0 to 10 min at 37°C. For DNA injection, incubations were extended 10 to 40 min. Aliquots were placed onto poly-L-lysine-coated slides and imaged using an Olympus IX83 inverted fluorescent microscope with a 60× oil immersion lens coupled to an Orca-Flash4.0 LT sCMOS camera and imaging system.

### DAPI fluorescent microscopy

To visualize nucleoids, DAPI staining was used following an established protocol with minor changes (20). Briefly, aliquots from the overnight cultures were washed three times with PBS (pH 7.4). Then, a 10 µL aliquot of cells was spotted onto poly-L-lysine-coated slides and incubated for 10 min to allow cell attachment. Excess liquid was removed and the cells were rinsed with water. Next, 3 µL of Fluorogel-II with DAPI (Electron Microscopy Sciences, Hatfield, PA) was added to the attached cells. A coverslip was placed over the sample, and the cells were imaged at 100× magnification using a Nikon E800 fluorescent microscope equipped with a digital imaging system.

### Phage absorption

Phage adsorption was conducted following the method outlined in (7) with some modifications. Overnight cultures were washed twice in fresh BHI medium and adjusted to an OD_600_ of 1.0. Then, 0.9 mL of cells was mixed with 0.1 mL of phage (2 × 10⁸ pfu/mL) were mixed in BHI containing 10 mM CaCl₂ in a 1.7 mL Eppendorf tube. The tubes were incubated at 37°C for 40 min to allow phage adsorption. After incubation, the tubes were centrifuged at 8,000 rpm for five min. The positive control was the WT DPC6143 adjusted to the same OD_600_, while the negative control consisted of the same medium with phage but no cells.

Following centrifugation, supernatant aliquots containing unbound phage were serially diluted 10-fold and spotted on a lawn of DPC6143. Plates were incubated for one day, and plaques were enumerated. The number of plaques in the positive control was set as 100% phage adsorption, while the number of plaques in the negative control was set as 0% phage adsorption. The percentage of phage adsorbed for the various mutant strains was then calculated based on this comparison.

### Genomic sequencing and analysis

Chromosomal DNA extraction and purification were done using the Wizard Genomic DNA Purification Kit (Promega, Madison, WI, USA) with slight modifications. Prior to cell lysis, 4 mL of overnight culture was pelleted by centrifugation and resuspended in 1 mL of BHI medium. The cells were then incubated with 20 mg/mL lysozyme and 50 mM EDTA at 37 °C for one hour to weaken the cell wall. Subsequent extraction and purification followed the manufacturer’s protocol, and the DNA was rehydrated in sterile deionized water at 65 °C for one hr. DNA concentrations were measured using a NanoDrop spectrophotometer, and the DNA was stored at 4°C.

Whole-genome sequencing of WT *S. mutans* DPC6143 was done using a hybrid sequencing strategy combining Illumina and Nanopore technology (SeqCenter, Pittsburgh, PA, USA). Sequence were assembled and annotated using Unicycler 0.4.8 and Prokka 1.14.5, respectively. Sequence variant identification of resistant isolates was done with Illumina sequencing, where short reads were aligned against the reference genome to identify mutations.

Genomic analysis was done using the Integrated Microbial Genomes & Microbiomes database (21). For comparative genomics, Phylogenetic Profilers was used.

### Biofilm assay

Overnight cultures were back diluted 1:20 in fresh BHI medium and allowed to grow for 2-3 hr or until early log phase. Cells were further diluted to an OD_600_ of 0.005 and 200 μL aliquots were added to a 96-well plate. After a 24 hr at 37°C, non-adherent cells were removed, and wells were gently washed with PBS (pH 7.2). Adherent cells were then resuspended in 100 μL of PBS, and OD_595_ readings were obtained with an Accuris SmartReader^TM^ 96-T plate reader. The OD reading of WT strain was set as 100% biofilm formation. The percentage biofilm for each mutant was calculated as OD_mutant_ /OD_WT_ × 100. To control for cell growth, parallel wells of each strain were resuspended and OD_595_ determined.

## Results

### ɸAPCM01 host binding and DNA injection

ɸAPCM01 is a clinical isolate from saliva and is a lytic phage belonging to the Siphoviridae family with B1 morphology (4). Of 17 serotype c and e strains tested, only *S. mutans* DPC6143, a serotype e strain, was sensitive to ɸAPCM01. We obtained these reagents and confirmed that ɸAPCM01 forms clear plaques on DPC6143 lawns. To visualize phage-host interactions, we fluorescently labeled ɸAPCM01 DNA with SYBR Gold and observed its attachment to DPC6143 (Fig. 1). The phage exhibited a tendency to bind near division planes, though attachment was also observed in other regions. Shortly after phage addition (e.g., 5-10 min), SYBR Gold labeled DNA was observed entering cells. By 10 to 30 min post-infection, fluorescent DNA was predominantly localized inside cells, revealing successful injection of phage genomes.

**Figure 1.**
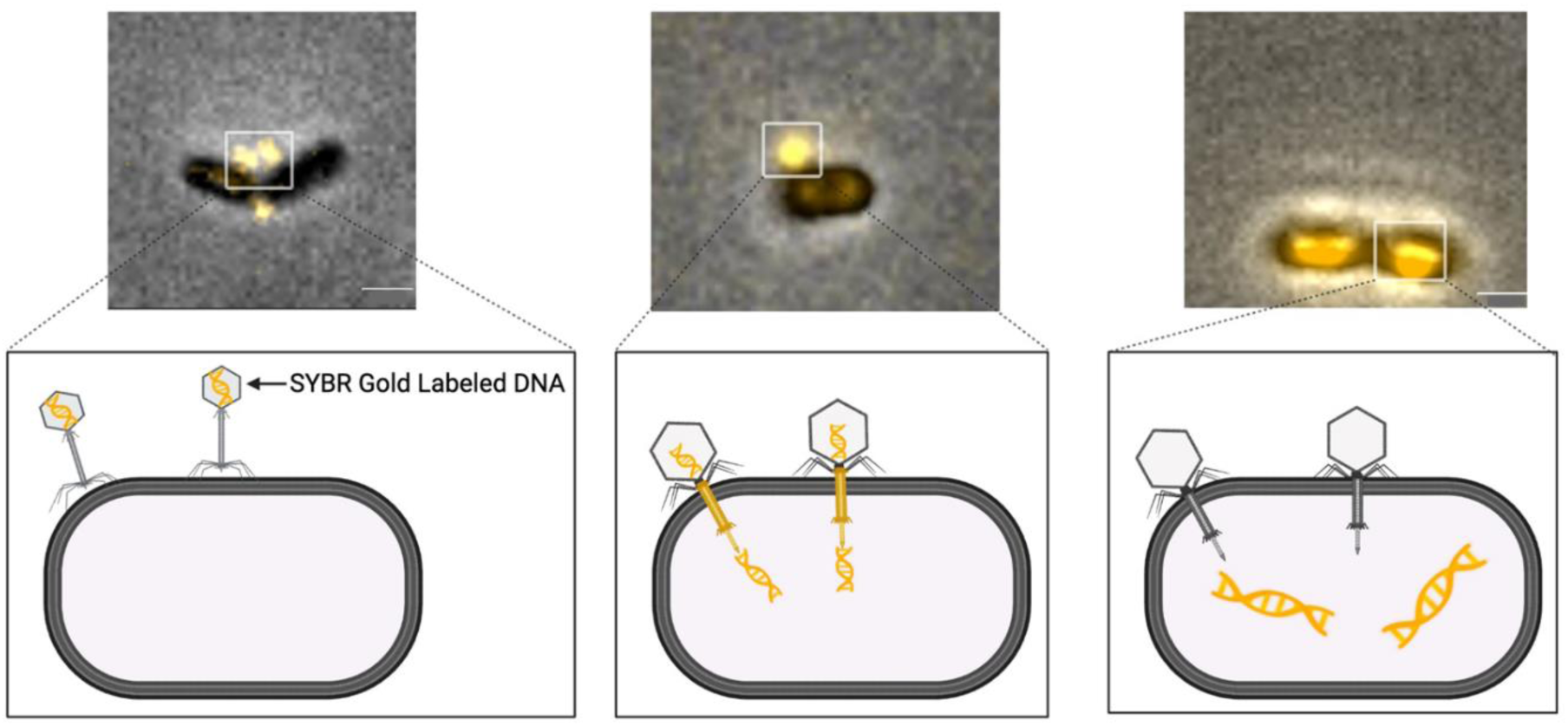
ɸAPCM01 binds to its host and injects its genome. The phage was labeled with SYBR Gold and incubated with DPC6143 host at different time points and stages of infection. Representative micrographs shown, and schematics illustrate different stages. Scale bars, 1 µm.

### DPC6143 genome sequence

To identify spontaneous ɸAPCM01-resistant mutations, we first generated a fully sequenced, assembled, and annotated reference genome for DPC6143 (NCBI accession number NZ_CP172847.1). The DPC6143 genome consists of a single circular chromosome comprised of 2,073,547 base pairs with a GC content of 36.90%, 65 tRNAs and five rRNA operons. Protein-coding sequences account for approximately 95% of the genes, organized into 1,982 predicted ORFs, which is larger than the model UA159-FR strain, which has 1,897 ORFs. Between these strains, 1,749 of the genes showed ≥ 95% identity, while 213 genes were unique to DPC6143 (≤ 30% identity).

DPC6143 contains a single CRISPR-Cas array that belongs to the type II-A Cas 9 system, which is found in other *S. mutans* strains such as UA159. The DPC6143 array contains 28 spacers (Table S1), several of which exhibit close sequence homology to phages M102, M102AD, SMHBZ8, and ɸAPCM01. Other spacers share homology with related phages identified from metagenomic sequences or *S. mutans* strain genomes. Although two of the spacers have 27/30 and 28/30 sequence identity to ɸAPCM01, strain DPC6143 was nevertheless sensitive to this phage.

### Isolation and sequencing of spontaneous phage-resistant mutants

To investigate phage resistance mechanisms, we isolated spontaneous mutants. *S. mutans* cultures were exposed to ɸAPCM01 in liquid media for 1 to 4 days. Aliquots from the lysates was then plated, and resulting isolated colonies were screened for resistance using an agar cross-streak method against ɸAPCM01. Validated clones were further tested for resistance by serial dilutions of phage on bacterial lawns. These screening steps were repeated multiple times with independent cultures to ensure no sibling isolates were obtained. In total 15 independent isolates were selected for genomic sequencing.

Table 1 summarizes the relevant mutations and ɸAPCM01 sensitivity of the isolates (Table S2 list all mutations). Notably, six out of 15 mutants contained an identical G382S substitution in RefSeq WP_002263319.1, a predicted glycosyl hydrolase 25 family member. Given that this identical mutation appeared in multiple unrelated isolates, we inferred it arose early in our frozen stock and was unrelated to resistance; thus it was excluded from further consideration.

**Table 1.** Summary of CRISPR spacers and relevant mutations in resistant isolates.

| Strains | Relevant mutations | pfu/mL |
| --- | --- | --- |
| DPC6143 | Wild type; clinical isolate | $1 \times 10^8$-$1 \times 10^9$ |
| *rgpX<sup>W190NS-CR1</sup>* | *rgpX* W190→NS CRISPR 1 | 0 |
| *rgpX<sup>P521H-CR2ab</sup>* | *rgpX* P521→H CRISPR 2a, 2b | $1 \times 10^{2*}$ |
| *rgpF<sup>E491K</sup>* | *rgpF* E491→K | $1 \times 10^{2*}$ |
| *rgpF<sup>T419P-CR3ab</sup>* | *rgpF* T419→P CRISPR 3a, 3b | $1 \times 10^{2*}$ |
| *rgpX<sup>FS</sup>* | *rgpX* ΔG codon 194 FS | $1 \times 10^2$-$1 \times 10^{3*}$ |
| *rgpF<sup>G382S-A</sup>* | *rgpF* G382→S | $1 \times 10^{3*}$ |
| *rgpF<sup>G382S-B</sup>* | *rgpF* G382→S multiple *rRNA* mutations | $1 \times 10^{3**}$ |
| *rgpD<sup>F162Y</sup>* | *rgpD* F162→Y | $1 \times 10^{3*}$ |
| CR4 | CRISPR 4 | $1 \times 10^{3*}$ |
| CR5abc | CRISPR 5a, 5b, 5c | $1 \times 10^{4*}$ |
| CR6ab | CRISPR 6a, 6b | $1 \times 10^{4*}$ |
| CR7 | CRISPR 7 | $1 \times 10^{4*}$ |
| CR8ab | CRISPR 8a, 8b | $1 \times 10^{5*}$ |
| CR9abc | CRISPR 9a, 9b, 9c | $1 \times 10^{5*}$ |
| CR10 | CRISPR 10 | $1 \times 10^{6*}$ |

\*No individual plaques at that dilution, though faint clearing present. \*\*Mutant contains multiple *rRNA* mutations not present in other *RgpF* G382S-A strain. FS, frameshift.

### CRISPR spacer acquisition

Previous work have shown that *S. mutans* strains readily acquire new spacers in their CRISPR array when challenged with phage (13, 15, 22). Consistent with this, 10 of our sequenced isolates acquired at least one new CRISPR spacer with 100% identity (28-30 bp) to a segment of the ɸAPCM01 genome (Fig. 2, Table S3). In total, 18 unique spacers were identified, each targeted different regions of the ɸAPCM01 genome.

**Figure 2.**
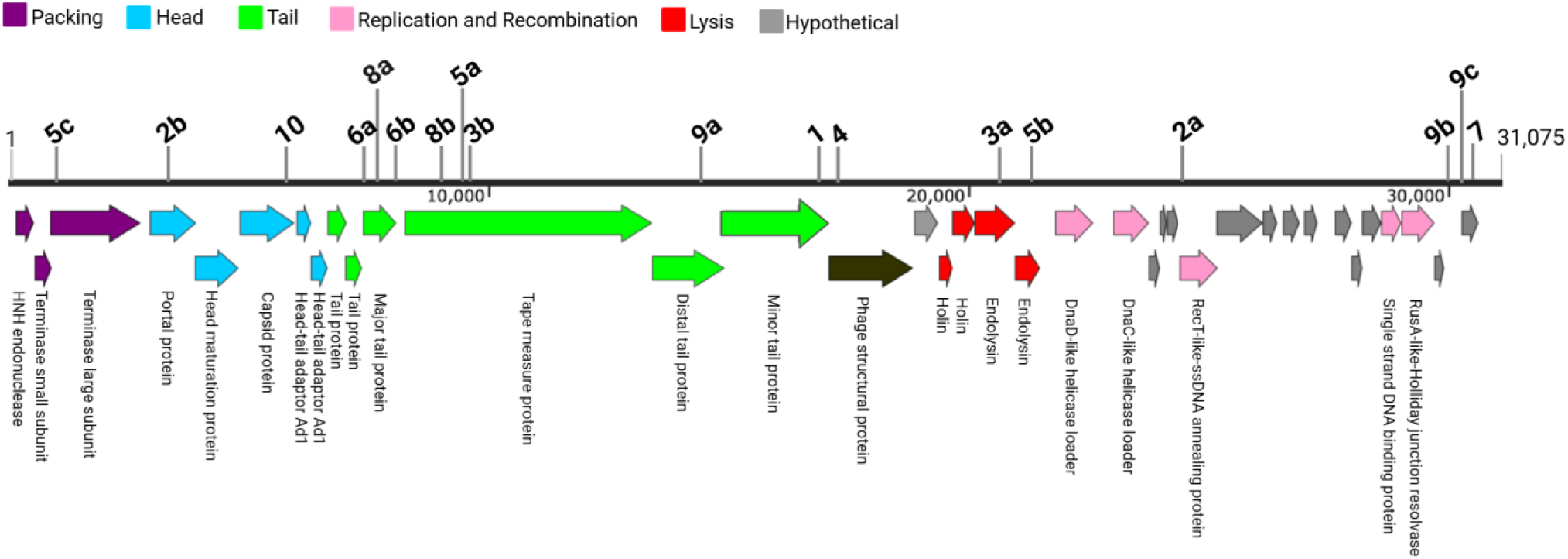
Genome organization of ɸAPCM01 and location of newly acquired spacers (4). Numbers with letters indicate multiple acquired spacers in isolate. See Tables 1 and S3 for details.

### RgpD, RgpF and RgpX mutants confer ɸAPCM01 resistance

Strikingly, eight of the 15 isolates had mutations in the *rgp* gene cluster, which is involved in rhamnose-glucose polysaccharide (RGP) biosynthesis (Table 1; Fig. 3). These mutations conferred the highest level of resistance to ɸAPCM01. Among these eight *rgp* mutants, three also harbored one or more acquired CRISPR spacers. Our screen identified one *rgpD* missense mutation, four *rgpF* missense mutations (two of which were identical in independent isolates), and three in a gene we named *rgpX*. Of the *rpgX* mutations, two were nonsense (NS) or frameshift (FS) mutations, likely resulting in null alleles (Tables 1 and S2). Notably, the missense mutations in *rgpD*, *rgpF* and *rgpX* affected highly conserved residues across *Streptococcus* species (Figs. S1 and S2) (23). The locations of all amino acid substitutions are illustrated on AlphaFold-predicted structures of these three proteins, where RgpF substitutions clustered within the same 3D region (Fig. S3) (24).

**Figure 3.**
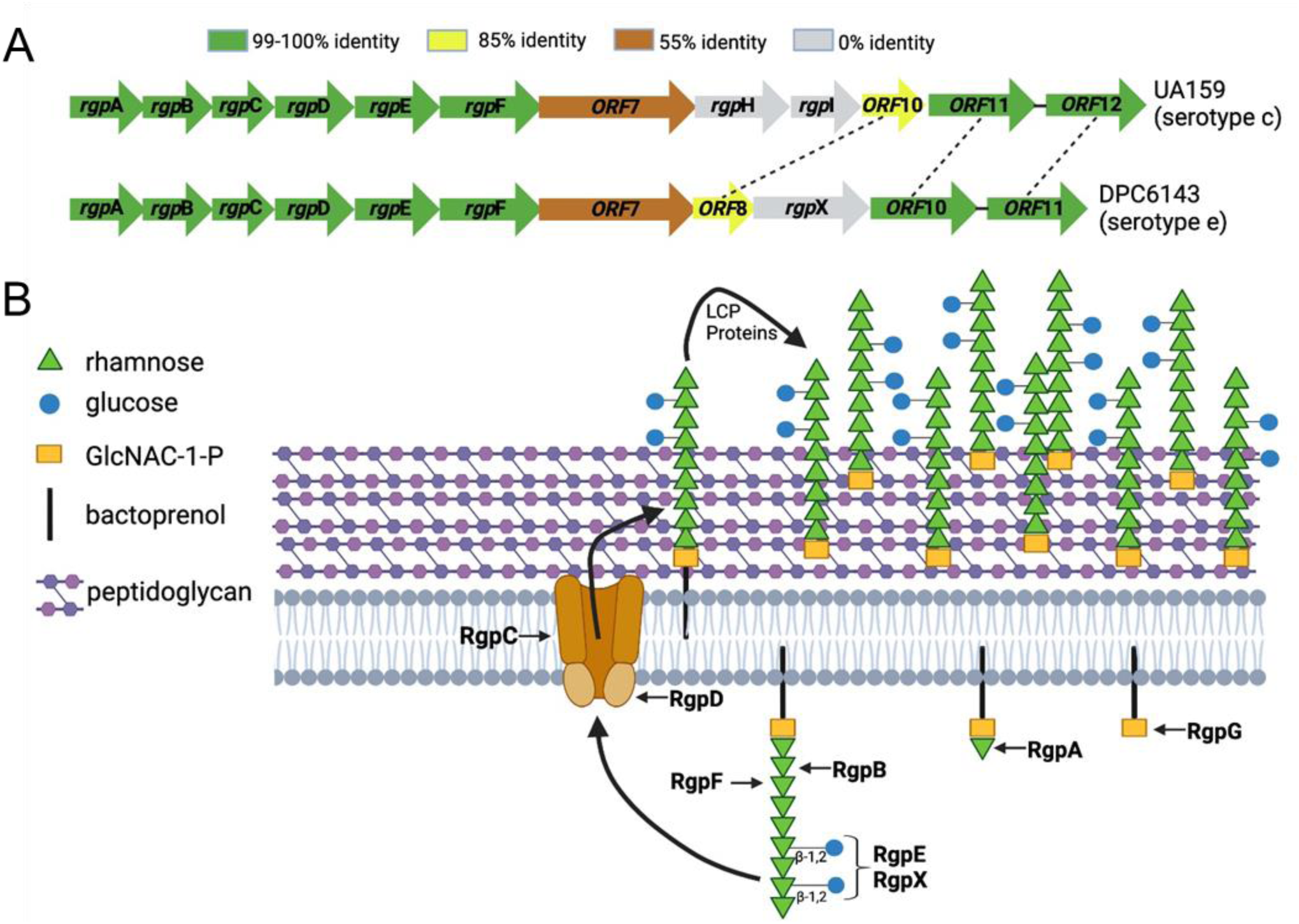
Gene organization and rhamnose-glucose polysaccharide (RGP) biosynthetic pathway. A) Comparison of *rgp* gene clusters from two serotype strains, showing percent protein identities. B) Proposed RGP biosynthetic pathway for serotype e, highlighting the hypothesized role of RgpX. Figure adapted from (25).

Attempts to construct complementing strains or knockout mutations in these genes were unsuccessful, as repeated attempts at electroporation and natural transformation with DPC6143 or its derivatives failed. Nevertheless, because we identified independent mutations in these genes, and in some cases in strain backgrounds free of other mutations, we conclude mutations in *rgpD, rgpF* and *rgpX* contribute to ɸAPCM01 resistance.

The functions of RgpD and RgpF have been previously characterized (25–28). RgpF acts as a rhamnosyltransferase responsible for elongating the rhamnose backbone of the polysaccharide, while RgpD serves as the ATPase subunit of an ABC transporter that translocates intracellularly synthesized RGP across the cytoplasmic membrane, where it is anchored to the cell wall (Fig. 3). These genes, along with the majority of the *rgp* gene cluster, exhibit close homology between UA159 and DPC6143.

In contrast, RgpX lacks a homolog in UA159 and remains uncharacterized. Sequence-based homology searches yielded a low-confidence score for a glycosylation enzyme family (E = 5×10^3^; COG1287), while DeepTMHMM 1.0 predicted RpgX contains 13 transmembrane helices (29). Additionally, structure-based searches using DALI revealed similarities to a mannose transferase (z-score = 13.7) (30). The *rgpX* gene is strategically positioned within the serotype-determining variable region of the *rgp* gene cluster (Fig. 3A), where it is unique to serotype e strains. Based on these findings, we propose that RgpX plays a role in modifying the side-chain of RGP. Specifically, RgpX may facilitate the formation of the β-1,2 linkages between rhamnosyl units and the glucose side chains, which are characteristic of serotype e.

### Rhamnose-glucose cell surface polysaccharides are essential for phage absorption

The serotype c RGP was previously described as the receptor for phage M102 (16). Building on this finding and our results, we hypothesized that RGP serves as a receptor for ɸAPCM01, a phage closely related to M102. To test this, we selected representative mutants for analysis; CR4 (CRISPR), *rgpD^F162Y^*, *rgpF^E491K^* and *rgpX^FS^*. These strains harbor either a single spacer or a single mutation in the respective genes, with no other mutations predicted to contribute to resistance (Table 1 and Fig. 4A).

**Figure 4.**
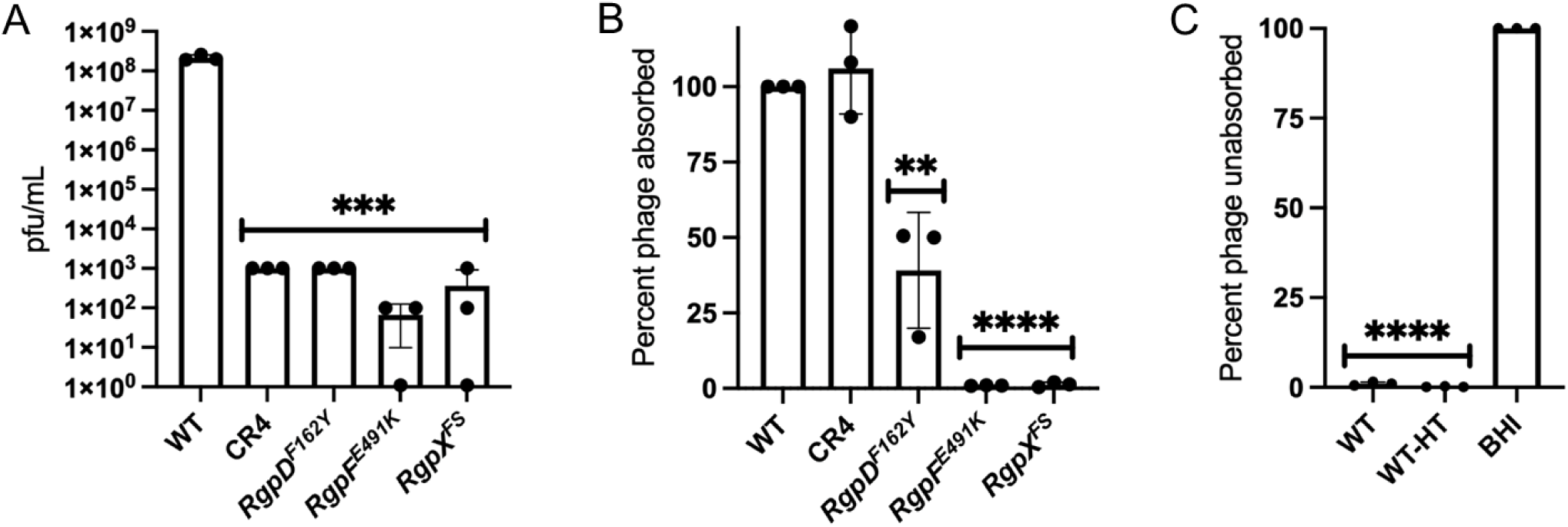
Resistance and absorption profiles to ɸAPCM01. A) EOP of isolates compared to DPC6143 (WT). B) Phage absorption of mutants compared to WT, set at 100%. C) Effect of 85 °C heat treatment (HT) on WT for phage absorption. BHI, media only. Assays were done in three biological replicas. Student t-tests; ** (p ≤ 0.01), *** (p ≤ 0.001), and **** (p ≤ 0.0001), significance compared to WT.

To assess binding, we used a phage adsorption assay. As shown in Figure 4B, phage adsorption was significantly reduced in the three *rgp* mutants, whereas CR4 exhibited no defect. Notably, among the *rgp* mutants, *rgpD^F162Y^* displayed a less severe defect compared to *rgpF^E491K^* and *rgpX^FS^*, suggesting that the F162Y substitution resulted in a partial loss-of-function mutation. This interpretation aligns with reports that *rgpD* null mutants are not viable (25, 27, 31).

To further investigate whether RGP serves as the receptor for ɸAPCM01, we tested phage adsorption following heat-treated, adapted a protocol previously used for phage M102AD (7). DPC6143 cells were incubated at 85°C for 30 min to denature proteins. Strikingly, despite the heat treatment, ɸAPCM01 continued to adsorbed efficiently to DPC6143 (Fig. 4C), supporting our conclusion that RGP, rather than a protein, serves as the receptor.

Previous studies reported that phage M102 and M102AD exhibit serotype c-specific binding and infections patterns (9, 16). To determine whether ɸAPCM01 was similarly restricted to serotype e, we tested UA159. As expected, ɸAPCM01 completely failed to form plaques or induce lysis on UA159 lawns (Fig. 5A). However, unexpectedly, the phage readily adsorbed to UA159 (Fig. 5B). To confirm this finding, we labeled ɸAPCM01 with SYBR Gold and visualized its interaction with UA159. The results showed that the phage successfully bound to UA159 and injected its DNA (Fig. 5C). These results suggest that UA159 resistance occurs after phage binding and DNA injection but before cell lysis.

**Figure 5.**
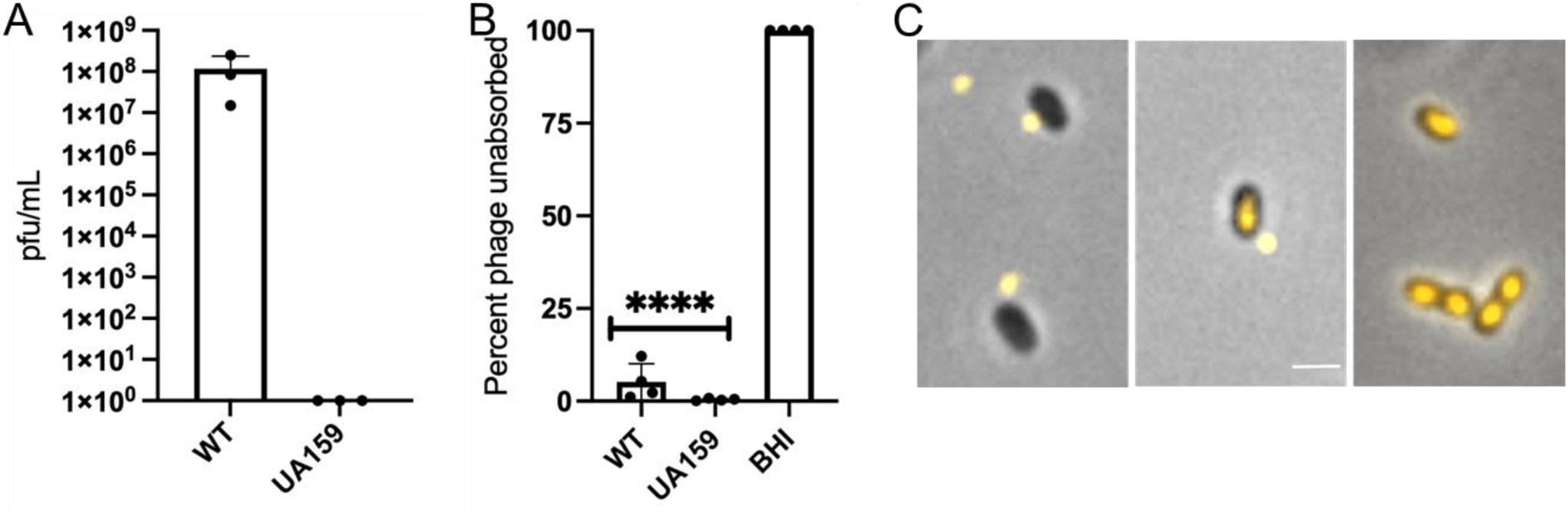
UA159 resistance and absorption to ɸAPCM01. A) EOP to ɸAPCM01. B) Strain absorption to ɸAPCM01. Assays were done in four biological replicates. Student t-tests; **** (p≤ 0.0001) significance compared to WT. C) Representative micrographs of SYBR Gold label ɸAPCM01 at different stages of binding through genome injection into UA159. Scale bar, 1 µm.

Although the mechanism of resistance remains unclear, UA159 contains two CRISPR-Cas systems with seven spacers, one of which matches ɸAPCM01 in 28 out of 30 bases. Taken together, these results indicate ɸAPCM01 binds to and injects its DNA into both serotype c and e strains, suggesting a broader host range than previously thought.

### Cell division defects in *rgpF* and *rgpX* mutants

Previous work demonstrated *rgp* mutants exhibit cell division defects (25, 26, 32). To assess these defects in our mutants, we stained nucleoids with DAPI and visualized cells using fluorescent microscopy. As shown in Figure 6A, the *rgpF^E491K^* and *rgpX^FS^* mutants displayed elongated chains and uneven daughter cell sizes, both hallmarks of cell division defects. In contrast, *rgpD^F162Y^* exhibited a morphology similar to WT.

**Figure 6.**
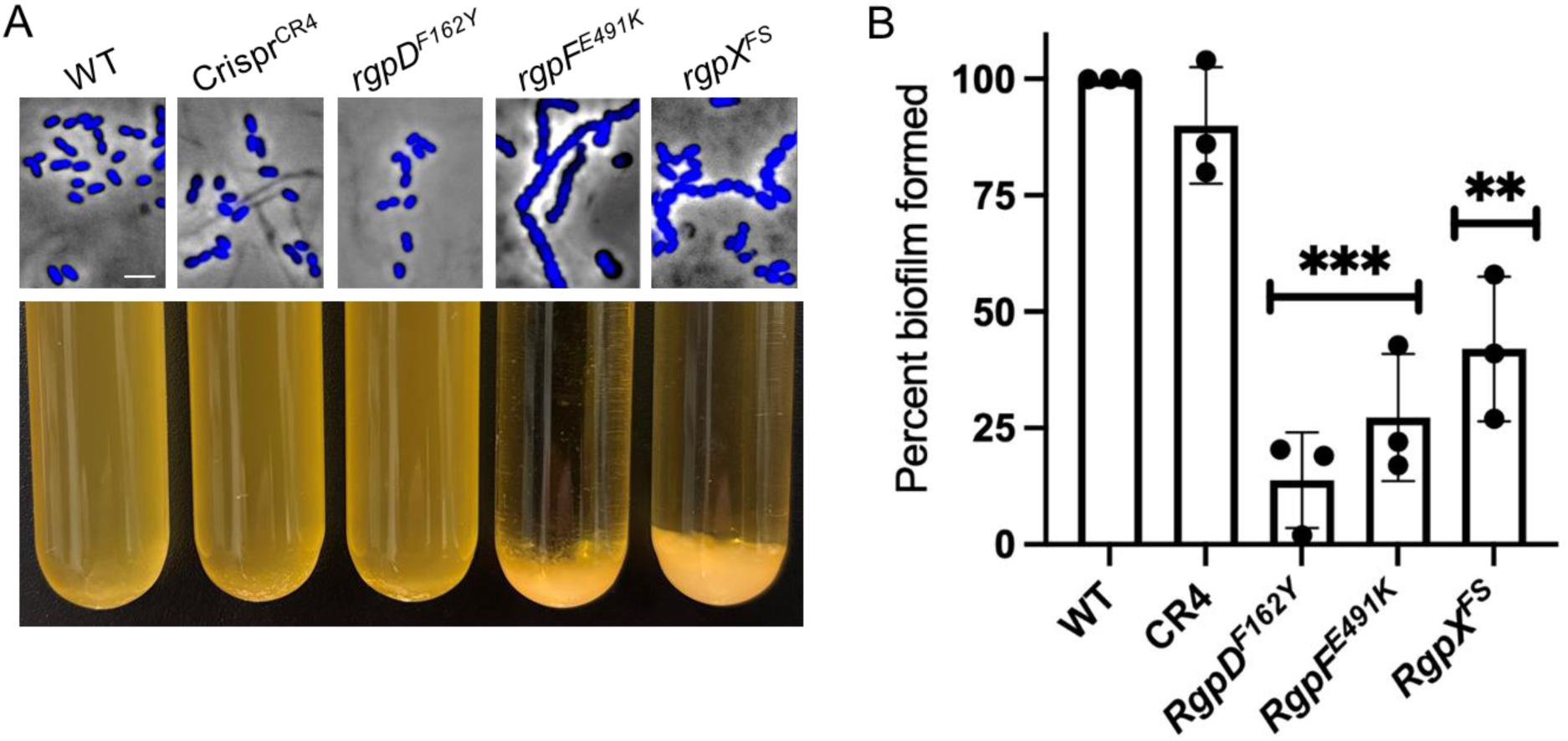
*rgp* mutant phenotypes. A) Representative morphologies of DAPI-stained strains (top). Sedimentation characteristics of overnight cultures (bottom). See Figure S4 for micrographs of sedimented *rgpF^E491K^* and *rgpX^FS^* mutants compared to WT. Scale bar, 2 µm. B) Biofilm formation after 24 hr incubation in microtiter wells. Following media removal and washing, biofilms were resuspended for OD_595_ measurements. WT set at 100% biofilm and compared to mutant values. Three biological replicates. Student t-tests; ** (p ≤ 0.01) and *** (p ≤ 0.001) significance values compared to WT.

Additionally, the *rgpF^E491K^* and *rgpX^FS^* mutants sedimented from overnight cultures−a phenotype not observed in *rgpD^F162Y^* or WT strains (Fig. 6A). To further investigate, we examined cells from the sedimented cells microscopically and found they formed tangled cells mats, likely explaining their sedimentation (Fig. S4). Finally, since *rgpD^F162Y^* did not exhibit a cell division or sedimentation defect and only had a limited ɸAPCM01absorption phenotype, this further supports the notion that it was a hypomorph allele.

### Biofilm defects of *rgp* mutants

To examine potential fitness trade-offs associated with phage resistance, we assessed biofilm formation in microtiter plate wells. Notably, the three *rgp* mutants displayed a reduced ability to form biofilms (Fig. 6B). Interestingly, *rgpD^F162Y^* exhibited the most severe biofilm defect. To control for cell growth, parallel wells were resuspended and OD_595_ readings showed that all cell culture densities were within 1-10% of each other. Notably, when sucrose was added to the assay, the biofilm defects were completely masked, suggesting sucrose induces an alternative biofilm-forming pathway (33, 34). These findings indicate that while *rgp* mutations confer phage resistance, they impose a fitness cost by impairing biofilm formation and, consequently, virulence.

## Discussion

Here, we characterized 15 spontaneous *S. mutans* isolates resistant to the lytic phage ɸAPCM01. Strikingly, 10 of these clones acquired one or more CRISPR spacers, generating a total of 18 new unique spacers that perfectly match ɸAPCM01 across various regions of its genome. Additionally, we sequenced the parent and clinical host strain DPC6143 and found it contained 28 original spacers, 11 of which matched different phages, with five being close matches to the M102, M102AD and ɸAPCM01 phage family. In conjunction with previous studies (13, 15, 22), our work further highlights that wild *S. mutans* strains express active CRISPR-Cas systems to combat phage attacks.

This is the first study to characterize spontaneous *S. mutans* phage resistance mutants beyond those acquiring new CRISPR spacers. Among the five resistant isolates that did not acquire spacers, they all had mutations in *rgpD*, *rgpF* or *rgpX*. Additionally, three other *rgpF* and *rgpX* mutants had acquired spacers. Phage absorption assays revealed that the *rgp* mutants − but not the CRISPR isolate − blocked ɸAPCM01 binding, indicating that the rhamnose-glucose cell surface polysaccharides serve as a phage receptor. This conclusion is supported by heat-treated cells retaining their ability to bind phage, as found in a prior study with M102AD (7).

The sole *rgpD* mutant contains an F162→Y substitution in a highly conserved residue. Previous research indicated that *rgpD* is an essential gene, and our phenotypic analysis suggests *rgpD^F162Y^* is a hypomorph allele (25). Curiously, all four of our *rgpF* mutants contain missense mutations that cluster in the same conserved region of the predicted structure. This implicates this region as essential for RgpF function and hints *rgpF* could be an essential gene in DPC6143. However, in UA159 Δ*rgpF* mutants were viable but exhibited significant fitness costs (25). Of the three *rgpX* mutants, two appear to be null alleles, suggesting that *rgpX* is not essential. Based on its chromosomal position within the *rgp* gene cluster and structural homologies, we hypothesize RgpX plays a role in catalyzing the β-1,2 linkages between rhamnosyl units and the glucose side chains, characteristic of serotype e strains. However, future studies are required to test this hypothesis.

Perhaps the most surprising finding is that phage ɸAPCM01 bound to and injects its DNA into UA159, a serotype c strain. This was unexpected because ɸAPCM01 and related *S. mutans* phages exhibit serotype specificity (4, 6, 9, 16). Our UA159 findings suggest that the side-chain linkages in both serotype c and serotype e are sufficient for ɸAPCM01 adsorption. In the literature, the clearest argument for serotype specificity comes from M102, which demonstrated absorption to all tested serotype c strains but not to serotypes e, f or k (16). These serotypes are distinguished by unique glucose side-chains linkages to the rhamnose backbone polymer (35). Notably, in the M102 study, the removal of serotype c-specific genes abolished absorption, while the introduction of serotype c-specific genes into sereotype e or f strains significantly increased M102 absorption, but not infection. In contrast, the recently isolated temperate phage ɸKSM96 infects both serotype c and e strains (8). Although a specific linkage between the glucose side chain and the rhamnosyl subunits does not appear to limit ɸAPCM01 host range absorption, the presence of a side chain still appears to be critical, as *rgpX* mutants block absorption. Our discovery that ɸAPCM01 binds to and injects DNA into a serotype c strain raises the possibility that it may infect certain serotype c strains. Why UA159 and other *S. mutans* strains are resistant to ɸAPCM01 remains to be elucidated. Finally, SYBR Gold labeling provides a simple and useful assay to monitor phage bind and DNA injection, which can help unravel the stage at which host resistance occurs.

Although mutations in *rgp* genes confer resistance to ɸAPCM01 and other *S. mutans* phages (16), they come with fitness costs (25, 26, 32). Specifically, *rgpD*, *rgpF*, and *rgpX* mutants are defective in biofilm formation, while *rgpF* and *rgpX* mutants also exhibit severe cell division abnormalities. These findings align with previous reports revealing that *rgp* mutants have multiple defects, including impairments in general stress protection and virulence, ultimately compromising their pathogenic potential.

The cell division defects in *rgp* mutants are linked to GbpB, a presumed cell wall hydrolase responsible for proper peptidoglycan degradation at the division site (36, 37). Here, GbpB relies on the RGP complex for correct localization, and its mislocalization results in abnormal cell morphology characterized by elongated chains and unevenly sized daughter cells.

Finally, to develop an effective therapy to treat dental caries, a phage cocktail will most likely be required. This strategy helps to overcome limitations of narrow serotype and strain specificity found in many *S. mutans* phages and provides a means to combat host resistance mechanisms. Given our and prior findings, *S. mutans* strains broadly express active CRISPR-Cas systems that generate adaptive immunity to phage. To overcome this challenge, *S. mutans* phages can potentially be engineered with anti-CRISPR systems, such as the AcrIIA6 system found in *S. thermophilus* phage genomes (38).

## Abbreviations

RGP: rhamnose-glucose polysaccharide
BHI: Brain Heart Infusion
EOP: efficiency of plating
PFU: plaque forming units
CRISPR: clustered regularly interspaced short palindromic repeats

## Acknowledgements

Phage ɸAPCM01 and its host strain were kindly provided and by Dr. Colin Hill. We thank Dr. Jacqueline Abranches for helpful suggestions and reagents. We thank Dr. Tingting Guo, Dr. Mike Weltzer and Pravas Roy for technical assistance.

## Supplemental Material

**Table S1.**
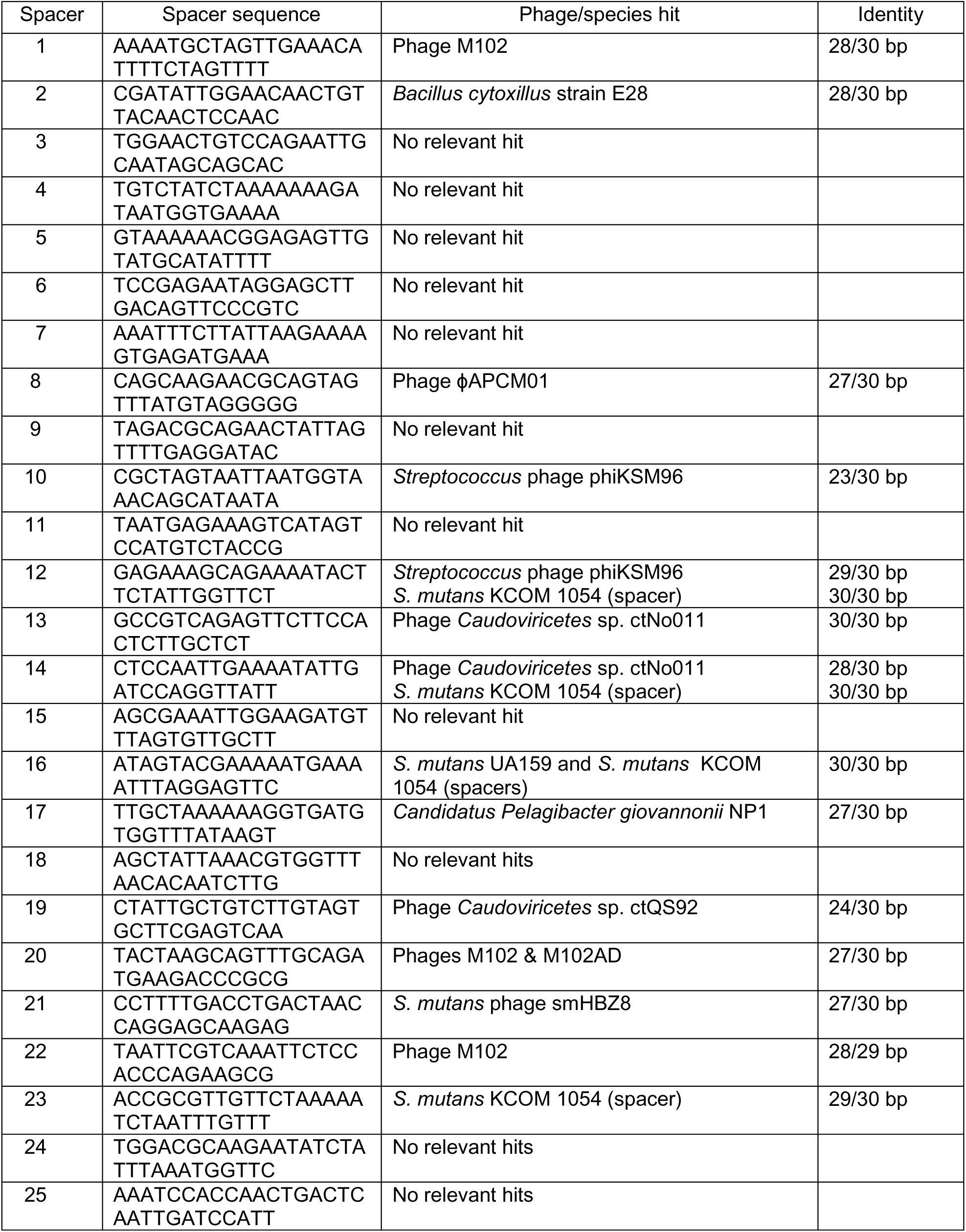
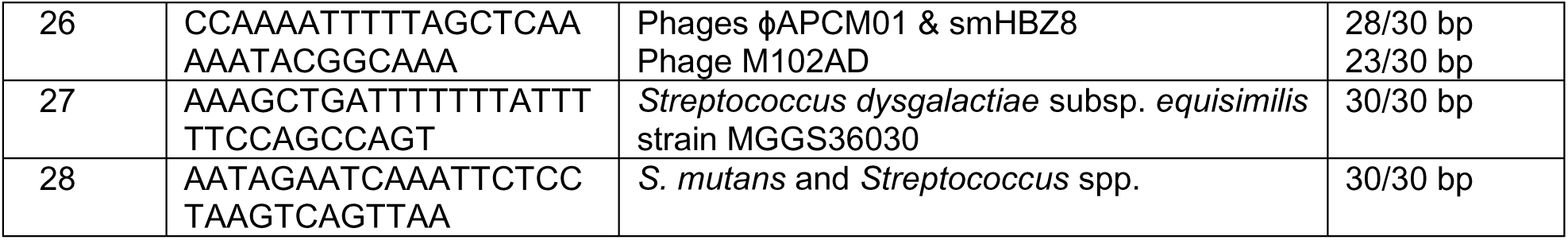
Endogenous CRISPR spacers in host DPC6143.

**Table S2.**
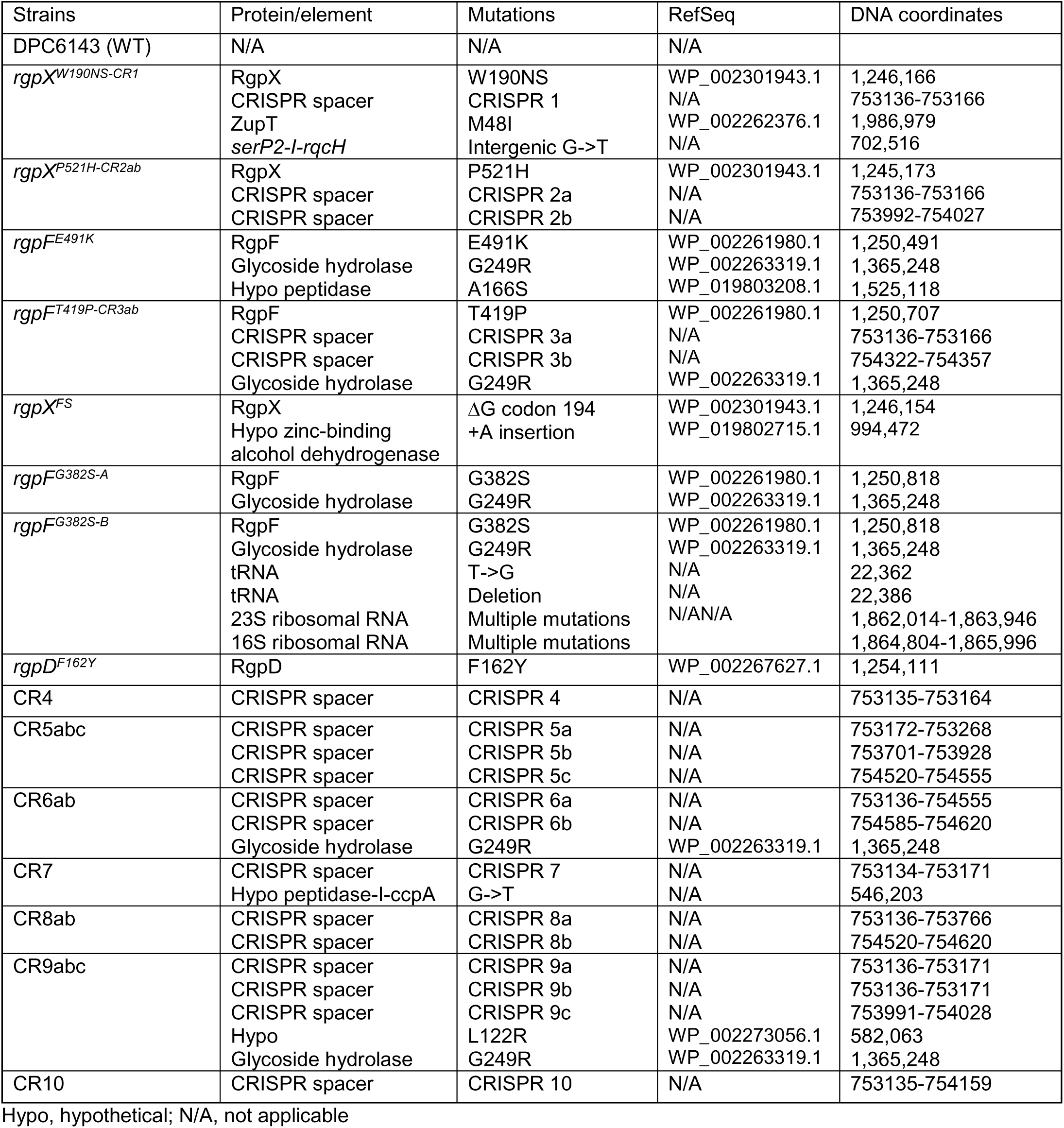
Mutations in ɸAPCM01 resistant isolates.

**Table S3.**
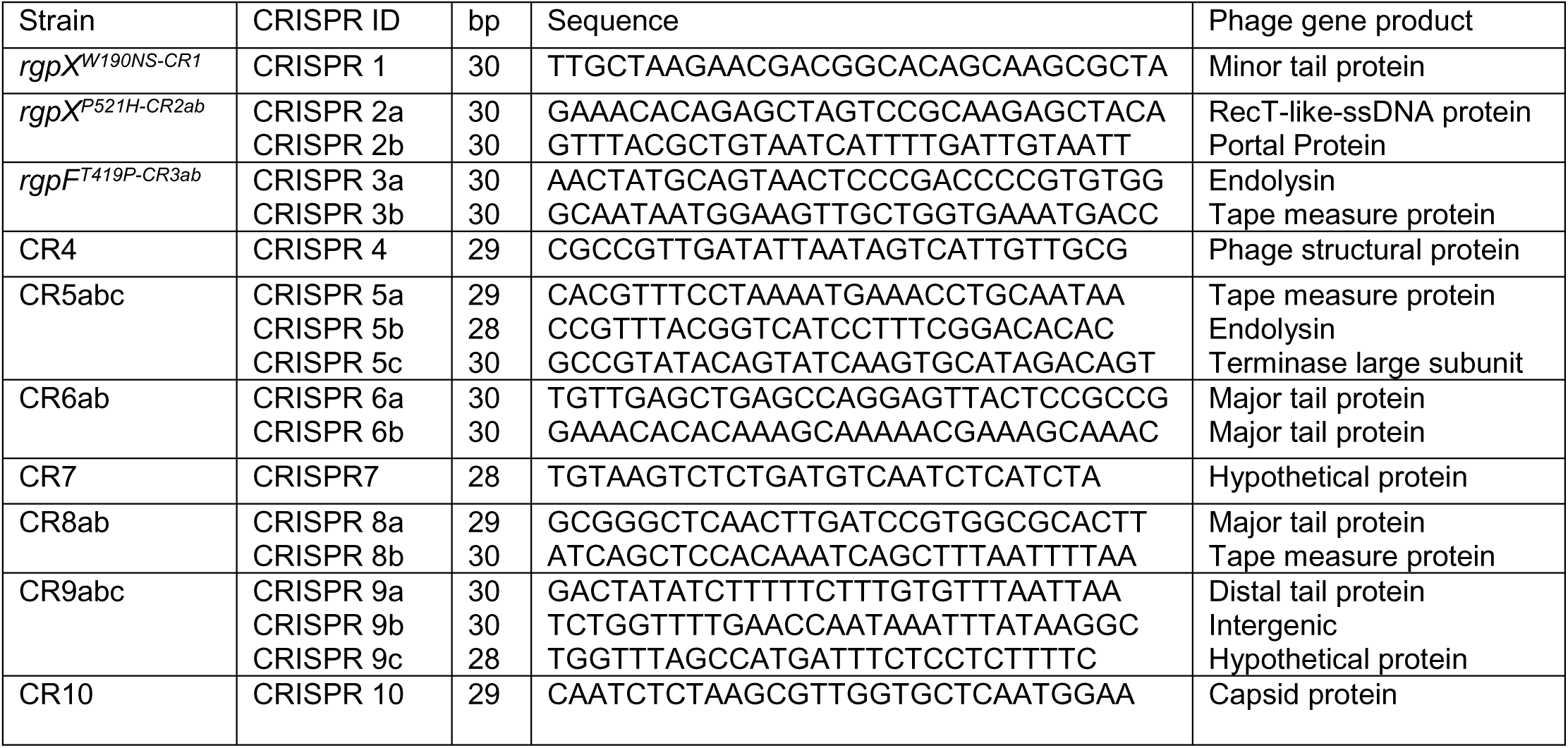
DNA sequence of acquired spacers and ɸAPCM01 targets.

**Figure S1.**
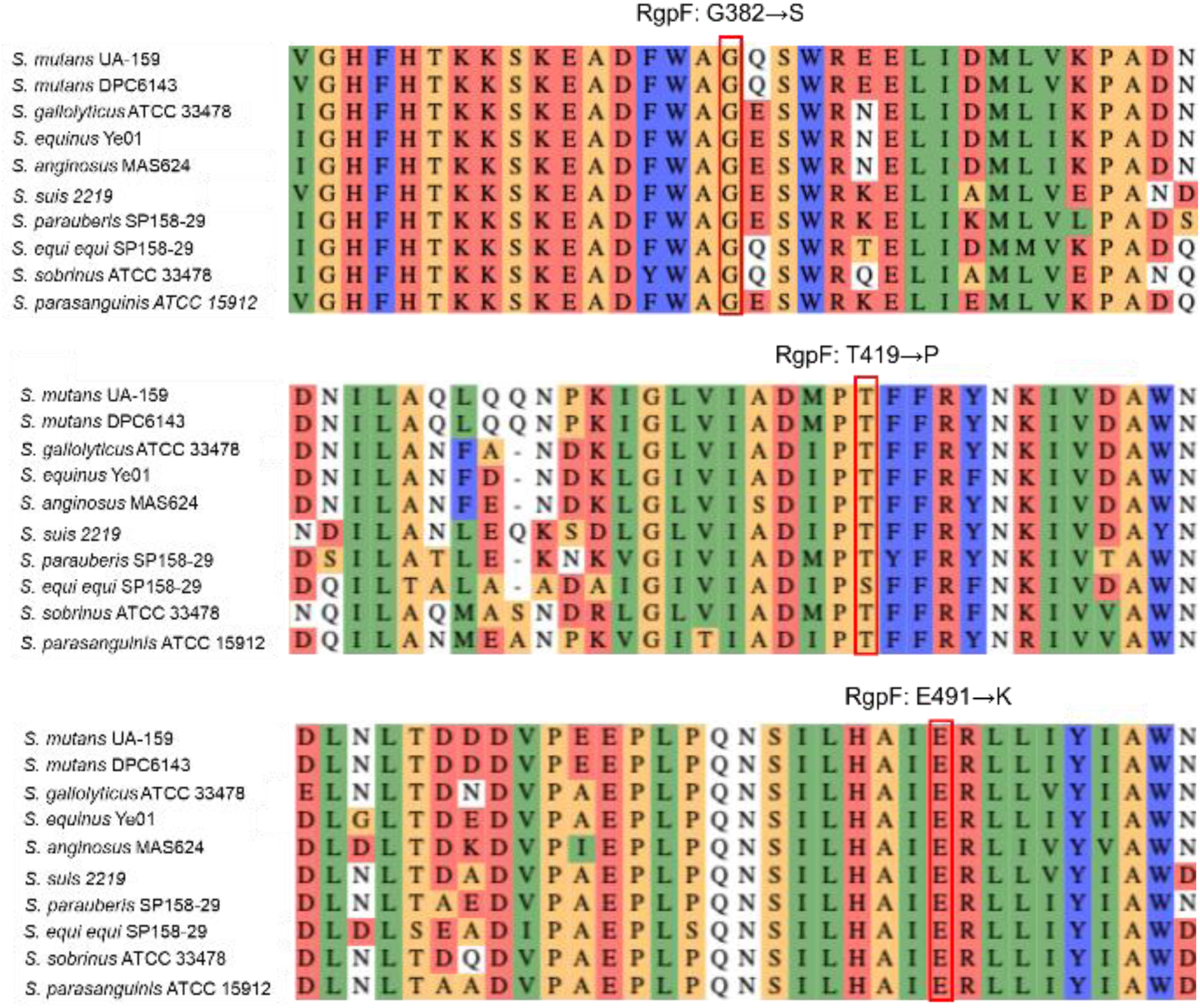
Sequence conservation among *Streptococcus* species around *S. mutans* RgpF substituted residues that confer ɸAPCM01 resistance.

**Figure S2.**
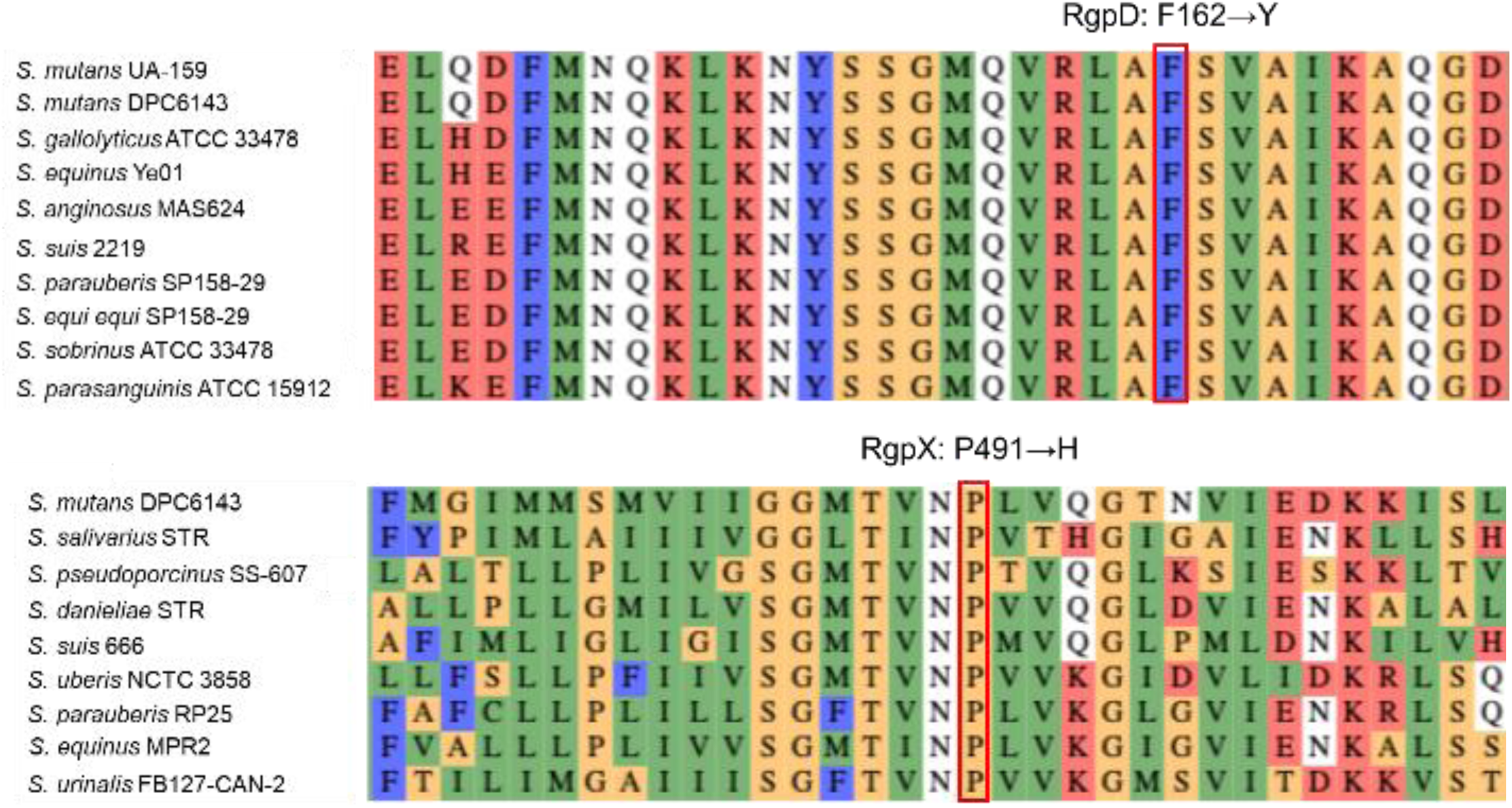
Sequence conservation among *Streptococcus* species around *S. mutans* RgpD and RgpX substituted residues that confer ɸAPCM01 resistance.

**Figure S3.**
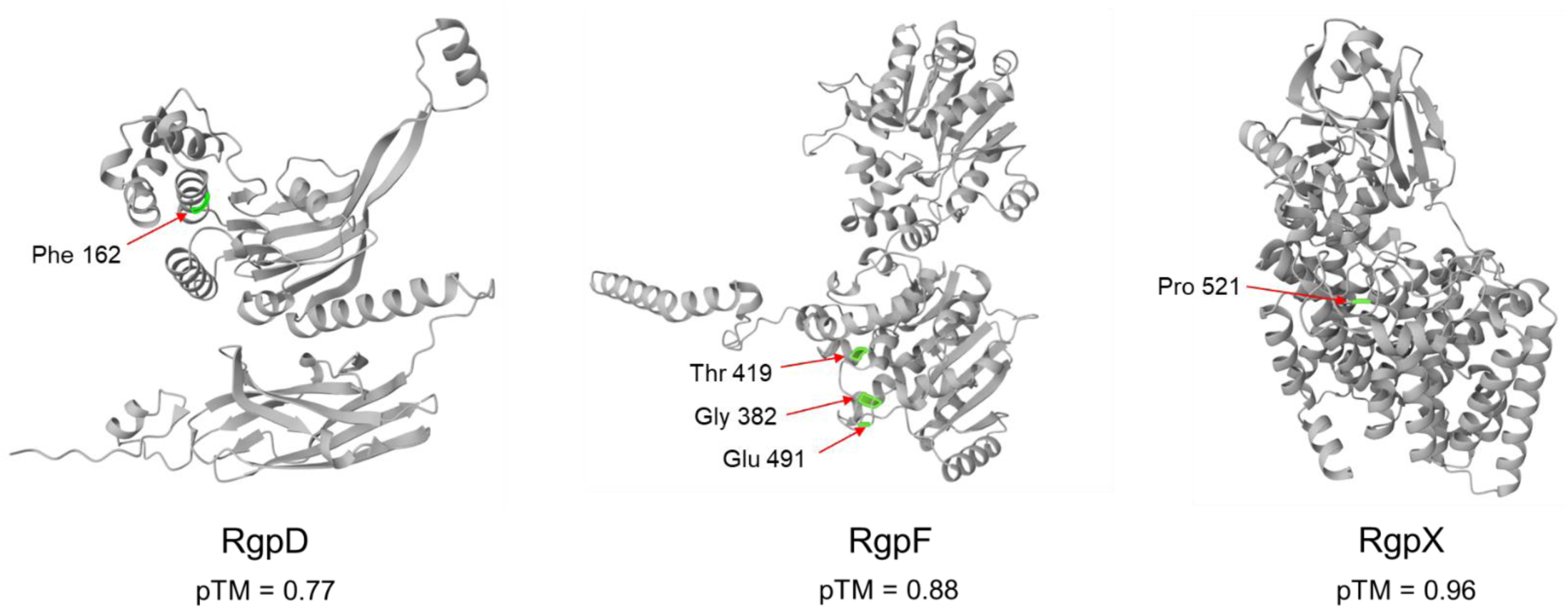
AlphaFold 3 predicted structures of Rgp proteins. Residues with substitutions that confer ɸAPCM01 resistance highlighted. All pTM scores indicate high confidence structures.

**Figure S4.**
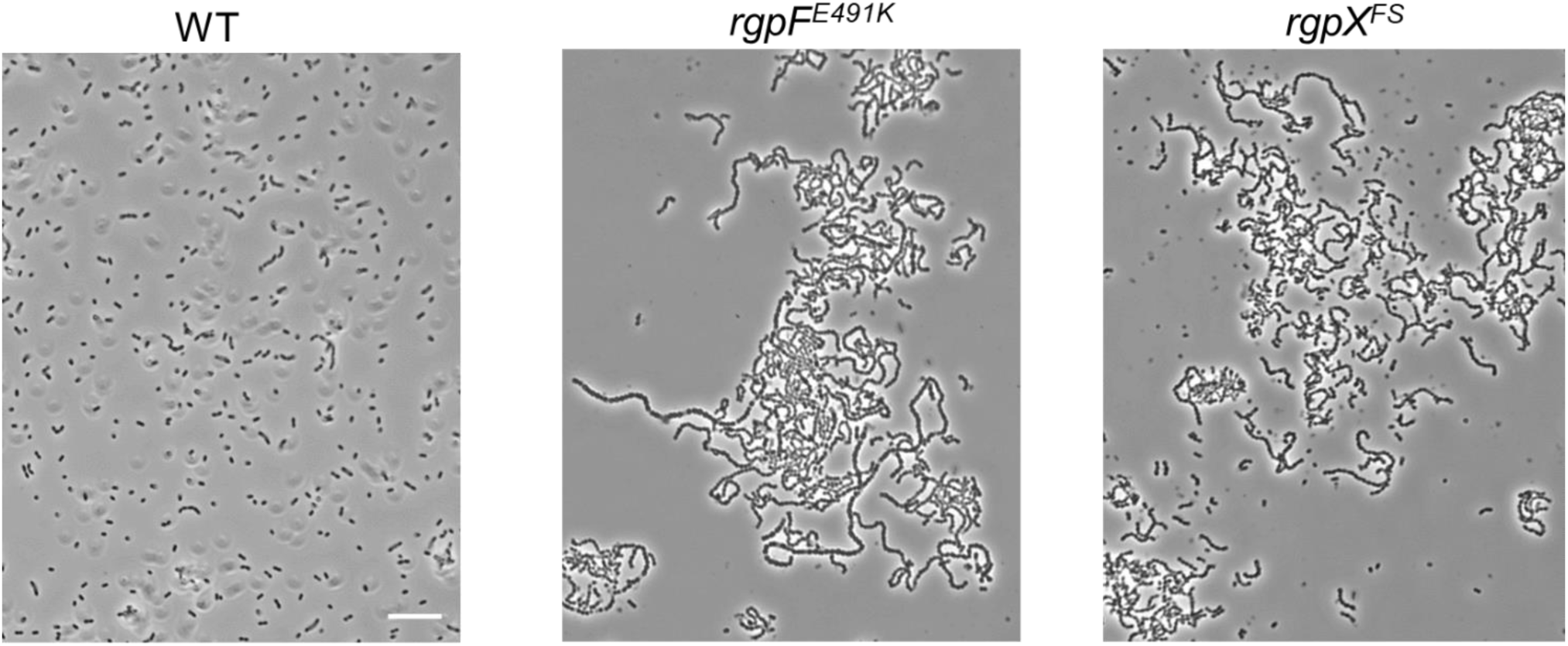
Culture tube samples from sediment shown in Figure 6A. Scale bar, 10 µM.

